# Top-Down Individual Ion Mass Spectrometry Reveals 85-110 kDa Catenin Phospho-Proteoforms Regulated by Actomyosin Contractility

**DOI:** 10.1101/2025.09.06.674621

**Authors:** Che-Fan Huang, Taojunfeng Su, Annette S. Flozak, Cara J. Gottardi, Neil L. Kelleher

**Affiliations:** Departments of Chemistry, Molecular Biosciences, and Proteomics Center of Excellence, Northwestern University, 2170 Campus Dr. Silverman B550, Evanston, Illinois 60208, United States; Departments of Medicine (Pulmonary and Critical Care), and Cell & Developmental Biology, Feinberg School of Medicine, Northwestern University, 303 E. Superior St. Simpson-Querrey 525, Chicago, Illinois 60611, United States

**Keywords:** Catenins, Phosphorylation, Proteoform, Mass Spectrometry, Top-Down Proteomics

## Abstract

A central challenge in top-down proteomics is the characterization of large proteoforms (>70 kDa) due to their high spectral complexity in mass spectrometers. Here, we advance individual ion mass spectrometry (I^2^MS) for intact mass and fragmentation analysis of β- and α-catenins (85-110 kDa), key components of adherens junctions. Using denatured I^2^MS, we resolved discrete phosphorylation states of catenins isolated from HEK cells subjected to differential actomyosin tension. Up to 10 phosphorylations were detected on β-catenin and 7 on α-catenin, with site-specific changes corresponding to actomyosin contractility. Notably, phosphorylation at α-catenin S641 was constitutive, while other sites in the P-linker and actin-binding domains as well as β-catenin S675 and S552 were sensitive to actomyosin perturbation. Application of I^2^MS for fragment ion detection (I^2^MS^2^) also enabled 25-30% sequence coverage for these exceptionally large proteoforms, compared to <1% using conventional methods for top-down mass spectrometry. Our results support a “catenin phospho-code” model, wherein combinatorial phosphorylation patterns encode mechano-transductive signals regulating cell–cell adhesion. This work establishes top-down I^2^MS as a viable approach for probing complex post-translational modification landscapes in high-mass proteins and highlights proteoforms as functional units in cellular regulation.

## Introduction

Protein post-translational modifications (PTMs) play critical roles in cellular signaling and the regulation of protein function. Often, multiple modification sites act in concert to enable higher-order or hierarchical control mechanisms (e.g. the histone code).^[1-5]^ As a result, the concept of the proteoform—defined as molecular variants of proteins arising from a single gene—has emerged as the new currency of proteomics research.^[6-8]^ Proteoforms capture the exact primary structure of individual protein molecules, providing a more functionally relevant and disease-associated perspective than analyses based solely on genes, protein groups, or tryptic peptides.^[9-12]^ Traditional proteomics methods, particularly bottom-up proteomics (BUP), infer protein identity by mapping tryptic peptides to protein groups. However, this approach is inherently limited for proteoform-level analyses due to the so-called “inference problem”.^[13]^ In contrast, top-down proteomics (TDP) circumvents this limitation by directly analyzing intact proteins through mass spectrometry (MS), thereby providing a more accurate framework for the characterization of proteoforms.^[14]^

A significant challenge in TDP is the analysis of large proteins, primarily due to difficulties in chromatographic separation, limited isotopic resolution, and ion decay within mass spectrometers. Therefore, conventional liquid chromatography–mass spectrometry (LC-MS) workflows are typically restricted to proteins smaller than 30 kDa for top-down analyses.^[15-18]^ Recently, we introduced a transformative single-ion approach to intact protein analysis— individual ion mass spectrometry (I^2^MS, **Figure 1**)—which enables isotopic resolution of proteins up to 466 kDa under native conditions.^[19-20]^ This technology presents new opportunities for directly profiling proteoforms exceeding 30 kDa. However, under commonly used denaturing electrospray conditions, such as formic acid buffers that promote higher charge states and minimize protein–protein interactions, I^2^MS remains limited to intact proteins below 70 kDa.^[21-22]^ Additionally, its application to MS^2^-level fragmentation analysis (I^2^MS^2^) has so far been demonstrated only for proteins under 45 kDa, including MEK1 and Tau.^[23-24]^ In this study, we aim to extend further the utility of I^2^MS for intact mass analysis of proteins >100 kDa and evaluate its performance in detecting fragment ions from large precursors such as catenins—a set of adhesive 85-110 kDa proteins found at the adherens junctions.

**Figure 1.**
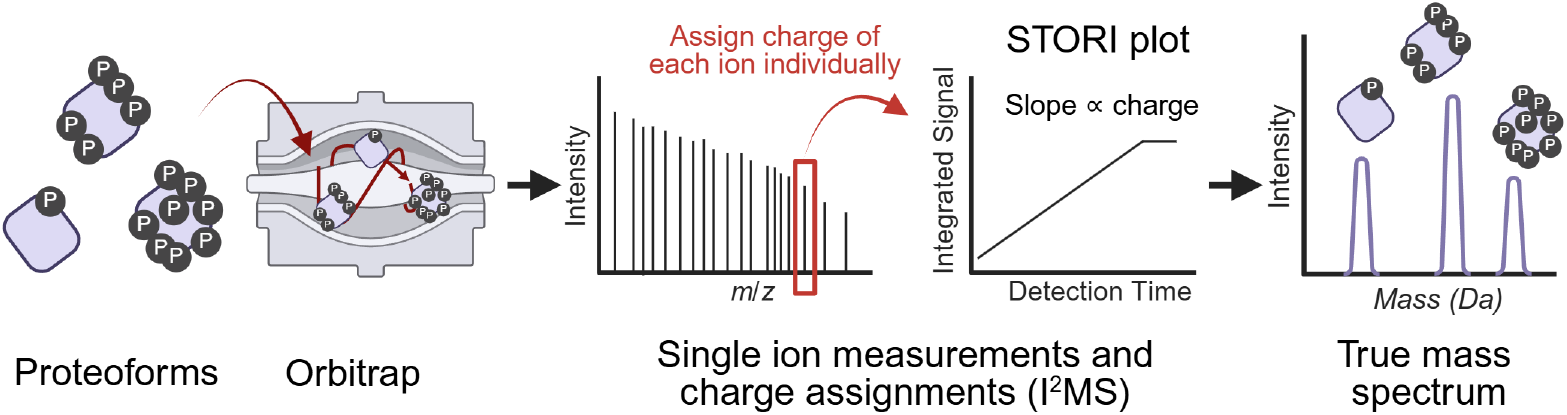
The process of Individual ion mass spectrometry (I^2^MS). Proteoform ion charge can be determined by monitoring its signal accumulation in orbitrap, resulting in a mass spectrum that reduces the complexity and improves detection versus traditional *m/z*-domain mass spectra.

Cell-cell cohesion is essential to the organization of multi-cellular tissues. One type of adhesive structure is the adherens junction, which in epithelial cells comprises transmembrane E-cadherin (E-cad) that recognizes the same protein on adjacent cells together with associated β-catenin (β-cat) and α-catenin (α-cat) to stabilize and link cadherins to the underlying actin cytoskeleton (**Figure 2a**).^[25-27]^ While the requirement of each component to tissue organization has been established, we know little about PTMs and structural changes that regulate cadherin-catenin-based adhesions associated with dynamic tissue states. Addressing this knowledge gap is important, as proper regulation is essential to important morphogenetic events such as mitosis and epithelial sheet invaginations.^[28-29]^

**Figure 2.**
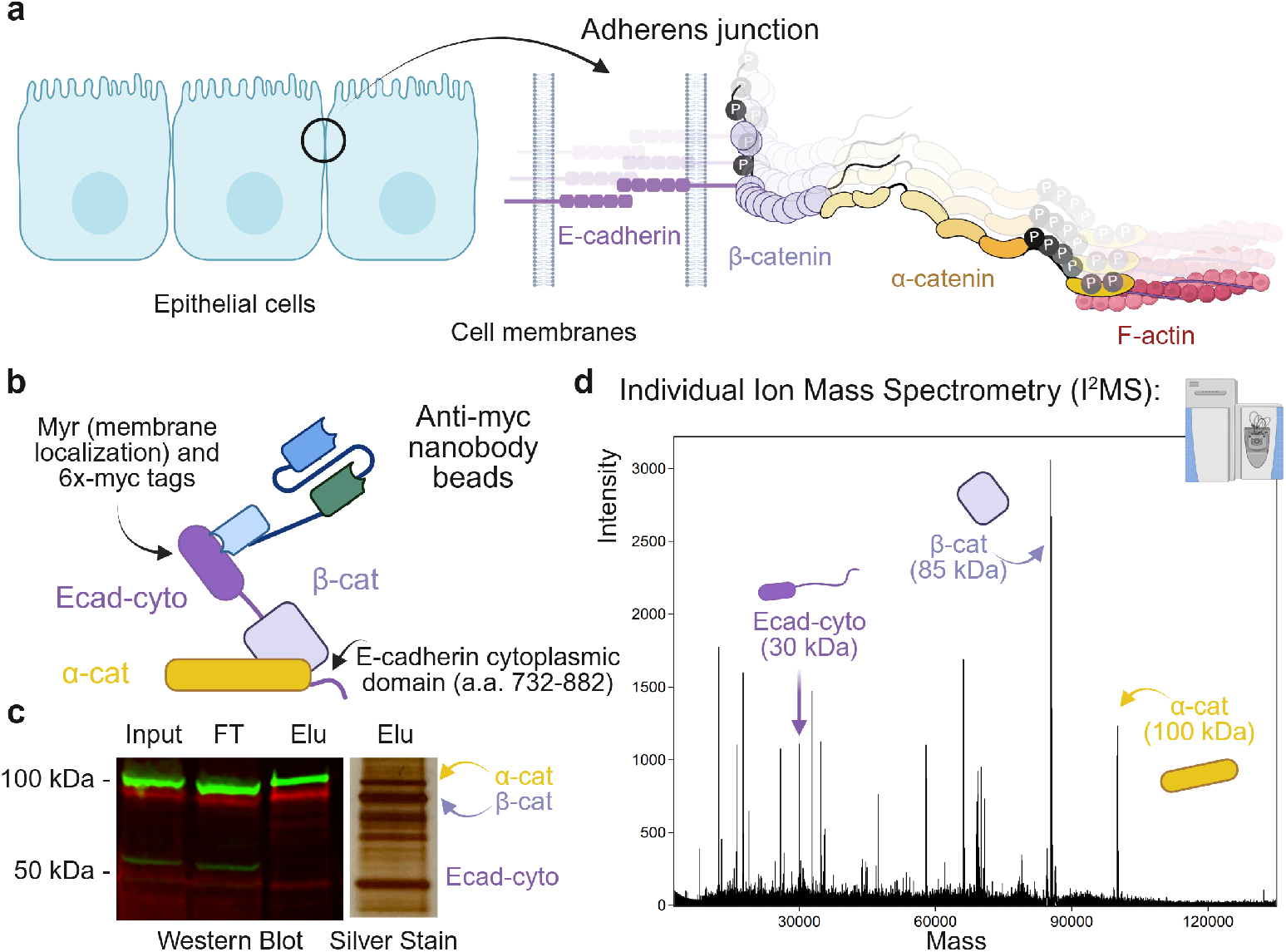
Isolation of the cadherin-catenin complex for mass spectrometric analysis. (a) Adherens junction consists of E-cadherin, β-catenin and α-catenin that link to actin filaments. (b) Cadherin-associated catenins were isolated from HEK cells expressing a myristoylated (myr) membrane-targeted Ecad-cytoplasmic domain. (c) Western blot and silver staining confirmed the presence of the complex. (d) The eluate was analyzed using individual ion mass spectrometry for detection of intact masses. All three components were observed after enrichment and I^2^MS.

Phosphorylation appears to be a major post-translational modification regulating cell adhesion and cytoskeletal architecture.^[30-35]^ However, current understanding is largely limited to individual phosphorylation sites identified through antibody-based methods or BUP using tryptic peptides. Thus, the combinatorial phosphorylation state of proteins, and whether other chemically distinct modifications act in concert with phosphorylation to drive diverse adhesion signaling and biomechanical states remain unclear. Herein, we demonstrate that the cadherin-associated catenin phospho-proteoform landscapes are sensitive to treatments that perturb actomyosin contractility with cutting-edge denatured I^2^MS technology, suggesting a phospho-code hypothesis linking mechano-sensing catenin to their distinct phosphorylation patterns with dynamic tissue states.

## Results and Discussion

### Optimization of Denatured I^2^MS for >100 kDa Proteoforms

To extend the capabilities of I^2^MS beyond the current ∼50 kDa upper mass limit,^[21-22]^ we first optimized the method using a GST-fused recombinant human β-catenin expressed in Baculovirus infected Sf9 cells (GST-β-cat, Abcam ab63175, ∼112 kDa). Ion injection time was controlled by Automatic Ion Control (AIC) to ensure optimal ion density for detecting single-ion events.^[36]^ Another critical acquisition parameter is the transient length—the duration over which the ion signal is acquired in the Orbitrap analyzer. In conventional I^2^MS, transient lengths typically range from 1 to 2 seconds. Previous studies have demonstrated that ultralong transients, extending up to 25 seconds, can substantially improve sensitivity and resolution in native I^2^MS applications.^[37]^ However, whereas native protein ions generally adopt compact, globular conformations that can persist within the mass spectrometer for extended durations, denatured protein ions exhibit larger collision cross-sections and are more prone to signal decay due to collisions and other destabilizing mechanisms.^[38-40]^ As a result, they are less suitable for ultralong transient acquisition. To determine the optimal conditions for analyzing denatured proteins >100 kDa, we evaluated spectra collected at 120k, 60k, 45k, and 30k resolution settings at 200 *m/z*, corresponding to transient lengths of 1.0, 0.5, 0.375, and 0.25 seconds, respectively, on a Q Exactive HF Orbitrap instrument. We found that a 0.375 s transient (45k resolution) offered the best balance, providing sufficient signal integration time to establish a defined slope in the STORI plot (**Figure 1**), while allowing for increased observation of individual ions within a fixed acquisition period—typically one hour for I^2^MS experiments. Data processing also plays a significant role in spectral quality. We employed an iterative voting algorithm for ion charge assignment and used the freely available STORIboard software for I^2^MS data analysis.^[22]^ For optimal performance at higher mass ranges with shorter transients, we adjusted several key processing parameters: lowering the minimum time-of-death threshold to 0.1 s to reflect the abbreviated transient; reducing the bin size from 5 ppm to 1.5 ppm to enhance mass resolution and accuracy; and setting the minimum ion count per bin to 1 to retain all detected ion events. A representative spectrum of GST-β-cat acquired/processed under these optimized conditions is shown in **Figure S1**, revealing at least five distinct proteoforms separated by 80 ± 1 Da, consistent with β-cat phosphorylation states—thus demonstrating I^2^MS sensitivity in denatured mode above the 100 kDa mass range.

### Isolation of Cadherin-Associated Catenins for Intact Mass Analysis

Catenins were initially identified through their interaction with cadherin-type cell–cell adhesion receptors, with decades of research underscoring their essential role in maintaining tissue cohesion and structural organization.^[25-27]^ In addition to their function within cadherin complexes, catenins also operate in alternative molecular contexts, where they contribute to diverse signaling pathways and cellular processes. For example, β-cat functions as a critical transcriptional co-activator in the Wnt signaling pathway^[41]^ while α-cat forms homodimers that can impact actin polymerization and filament organization.^[42-45]^ We hypothesize that the functional diversities of catenins are determined by their distinct proteoforms. Thus, it is important to isolate these proteoforms based on their specific cellular functions, rather than through indiscriminate global pull-down approaches. In this work, we focus on catenins relevant to cell-cell junctions (**Figure 2a**). To this end, we used a HEK cell line expressing the cytoplasmic domain of E-cadherin (Ecad-cyto), fused to a myristoylation (myr) sequence for membrane localization and a 6×Myc epitope tag to facilitate purification of cadherin–catenin complexes (**Figure 2b**).^[32]^ Western blot analysis detecting β-cat (green) and α-cat (red) confirmed that both proteins co-purified with Ecad-cyto as expected. Moreover, these catenins were the predominant proteins in the eluate, as observed by silver staining (**Figure 2c**).

The eluate was subsequently analyzed using I^2^MS for intact mass profiling. Proteoforms of Ecad-cyto (∼30 kDa), β-cat (∼85 kDa), and α-cat (∼100 kDa) were all detected (**Figure 2d**). Closer examination of β-cat and α-cat within the 85–110 kDa mass range revealed partially resolved isotopic patterns, enabling intact mass matching to their theoretical isotopic distributions (**Figures S2a** and **S3a**). Accounting for potential phosphorylation and oxidation events—possibly introduced during purification or electrospray ionization—we confirmed that cadherin-associated β-cat proteoforms exhibit N-terminal methionine removal and acetylation, along with zero to four phosphorylations under normal cellular conditions (**Figure S2b**). We note that there may be some phosphate adducts (+98 ± 1 Da) that were indistinguishable from a phosphorylation plus an oxidation (+96 ± 1 Da) and contributed to a positive shift in the observed molecular weight. Similarly, α-cat proteoforms consistently show N-terminal methionine removal and acetylation. Notably, a non-phosphorylated α-cat variant was not observed. Instead, α-cat phosphorylation ranged from one to three sites, suggesting a constitutive phosphorylation may be required for the formation of cadherin–catenin complexes (**Figure S3b**). Leveraging the ability of top-down mass spectrometry to survey the whole proteoform landscape, we determined that no other high-occupancy post-translational modifications (>5% abundance) were present, allowing us to rule out alternative modifications with confidence.

### Actomyosin Inhibitor/Activator Alters Catenin Phospho-Proteoform Landscapes

Increasing evidence suggests that both β-cat and α-cat are mechano-sensitive proteins, where enhancement of PTMs under force (e.g., membrane stretching) may impact cell-cell adhesion and signalling outcomes.^[46-47]^ To assess the effects of actomyosin contractility, HEK cells were treated with the actomyosin inhibitor blebbistatin, DMSO control, and the actomyosin activator calyculin A. Phenotypes were examined by fluorescent confocal microscopy following F-actin and DNA staining. Blebbistatin treatment (20 µM, 24 h) resulted in enlarged and flattened cells relative to the DMSO control, consistent with reduced actomyosin tension. In contrast, calyculin A treatment (20 nM, 2 h) led to actin retraction toward the nucleus and cell rounding, consistent with elevated actomyosin tension (**Figure 3a**). Furthermore, immunostaining with β-cat and α-cat antibodies (**Figure S4a**) revealed that both catenins increasingly enriched at cell–cell junctions in response to enhanced actomyosin activity (blebbistatin < DMSO < calyculin A, **Figure S4b**). We hypothesized that actomyosin force-dependent changes in catenin proteoform composition may contribute to adherens junction organization. Therefore, we next examined the proteoform landscapes of catenins under these treatment conditions.

**Figure 3.**
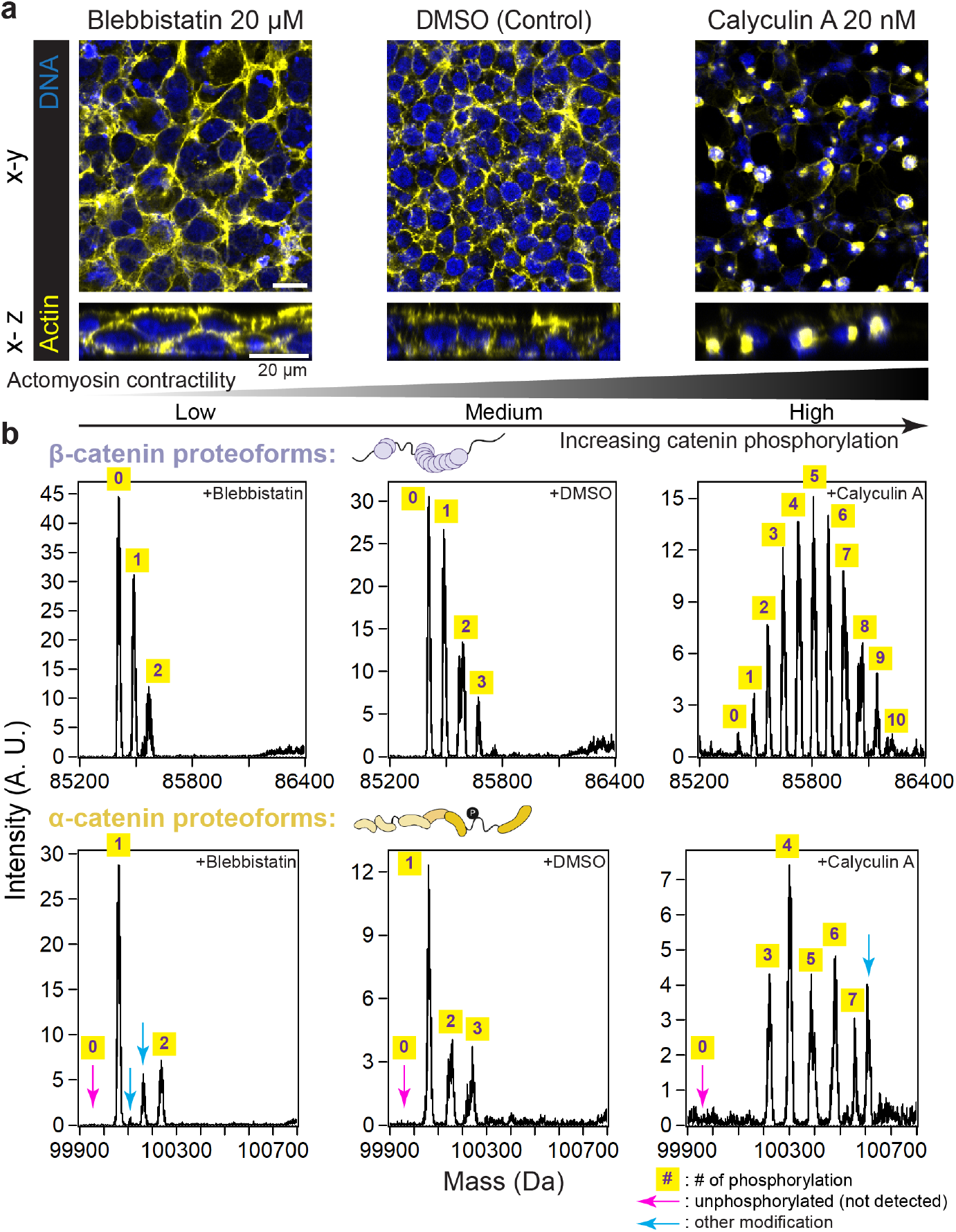
Intact masses reveal actomyosin-contractility-dependent modifications of cadherin cyto-domain associated catenins. (a) Confocal images of HEK cells under treatments that result in different levels of actomyosin contraction. F-actin is visualized with phalloidin (yellow); nuclei are stained with the DNA dye, Hoechst. Both *en face* (x-y; top) and orthogonal (x-z; bottom) views are shown. Scale bars: 20 µm. (b) β- and α-catenin proteoforms manifest different degrees of phosphorylation, where phospho-proteoform abundance is a function of actomyosin activity.

Cadherin-associated β-cat and α-cat were isolated from HEK cells treated with blebbistatin, DMSO, or calyculin A using the Ecad-cyto construct (**Figure 2b**), and their proteoform landscapes were profiled in triplicate using I^2^MS. Representative spectra spanning the corresponding proteoform mass ranges are shown in **Figure 3b**. The data revealed pronounced mass shifts consistent with phosphorylation (+80 ± 1 Da) as the primary PTM on both catenins. Notably, β-cat and α-cat responded similarly to treatments that modulate actomyosin contractility. The number and abundance of phosphorylation events decreased under reduced actomyosin activity (hypo-phosphorylation) and increased under elevated actomyosin activity (hyper-phosphorylation). Remarkably, we observed up to 10 phosphorylations on a single β-cat proteoform and up to 7 on α-cat. These proteoforms were clearly resolved in the mass domain—with isotopic resolution—thanks to the true mass detection enabled by I^2^MS. In contrast, under traditional ensemble MS, these proteoforms would have overlapped in the limited *m/z* space due to their high charge states (+80 to +120), which typically result in extensive spectral congestion (**Figure S5**).

Additionally, we made two notable observations regarding α-cat proteoforms. First, as noted above, α-cat phosphorylation ranged from one to three sites under normal cellular conditions. Here, we found that unphosphorylated α-cat was still absent from cadherin–catenin complexes purified from either blebbistatin- or calyculin A-treated cells (**Figure 3b**, pink arrows). This finding suggests that a constitutively phosphorylated site on α-cat may contribute to cadherin–catenin complex formation. Second, we observed mass shifts on α-cat that do not correspond to phosphorylation (**Figure 3b**, cyan arrows), suggesting that catenin phospho-states may be coordinately regulated by other PTMs. The observed shifts of +48 Da and +53 Da may reflect cysteine trioxidation or iron adducts—both commonly reported in MS experiments.^[48]^ Further investigation is needed to determine the specific sites and whether these modifications are endogenous.

### Identification of Mechano-Sensitive Catenin Phosphorylation Sites

Since phosphorylations were abundant in the calyculin A-treated cells, we used BUP to directly map catenin phosphosites without the phospho-peptide enrichment, typically required for phospho-peptide recovery.^[33]^ Ecad-cyto complexes purified from calyculin A-treated cells were digested with trypsin for LC-MS analysis and Resulting sequence coverage maps can be found in **Figure S6**. The observed phospho-peptides were further confirmed using MASCOT^[49]^ and FragPipe PTM analysis^[50]^ to distinguish between potentially ambiguous sites (**Table S1**). β-cat phosphosites found were: S29, S33, S37, T39, T41, T42, S45, S552 and S675. α-cat phosphosites were: S641, S652, T654, S655, T658, S851 and S886. All β-cat and α-cat sites identified are highly conserved across multiple cell lines and tissue types.^[33, 51-52]^ Two phosphosites in the actin-binding domain of α-cat (S851 and S886) were previously identified in large-scale phospho-proteomics data,^[53]^ although their function remains unknown.

Phospho-specific antibodies are available for most of the sites we identified, and we used them to assess the mechano-sensitive properties of each residue by comparing their intensities in the Ecad-cyto complexes purified from HEK cells treated under low (blebbistatin), medium (DMSO control), and high (calyculin A) actomyosin contractility conditions. We designate sites as actomyosin sensitive when they are both reduced by inhibition and enhanced by activation of actomyosin contractility via blebbistatin or calyculin A, respectively. Two β-cat residues were found to be mechano-sensitive: S675 and S552. pS675 intensity slightly increased (not statistically significant) as the tension increased from low to medium and showed a significant increase under high-tension conditions (**Figure 4a**). Similarly, pS552 intensity significantly increased with rising actomyosin contractility from low to medium and from medium to high (**Figure 4b**), suggesting that S552 is more mechano-sensitive than S675. This observation is consistent with a recent study by Fujita and coworkers, which identified S552 as the primary mechano-sensing site in fibroblast cell-sheet stretching.^[54]^ This supports our finding that S675 is less responsive to blebbistatin treatment. Whether there is a hierarchy between these two residues remains unclear. They may function independently, or pS675, together with mechano-force, may facilitate kinase access to S552, thereby enhancing its mechano-sensitivity. This will require further investigation at the proteoform level. The N-terminal phosphosites of β-cat are historically associated with the GSK3β-mediated degradation pathway of cadherin-free β-cat and are not considered mechano-sensitive.^[54-55]^ Western blots using three antibodies (targeting β-cat pS45, pT41/S45, and pS33/S37/T41, respectively) demonstrated that these sites were only detected in calyculin A-treated cells (**Figure S7**). While calyculin A induces substantial cell rounding and actomyosin contraction (**Figure 3a**) from inhibition of PP1/PP2A phosphatase activity and subsequent accumulation of phosphorylated myosin light chain,^[56]^ this observation may be explained by Ecad-cyto complexes being retracted from the membrane by actin filament, thereby activating the degradation pathway via N-terminal phosphorylations. However, we cannot rule out that the N-terminal phosphosites may also be PP1/PP2A substrates.

**Figure 4.**
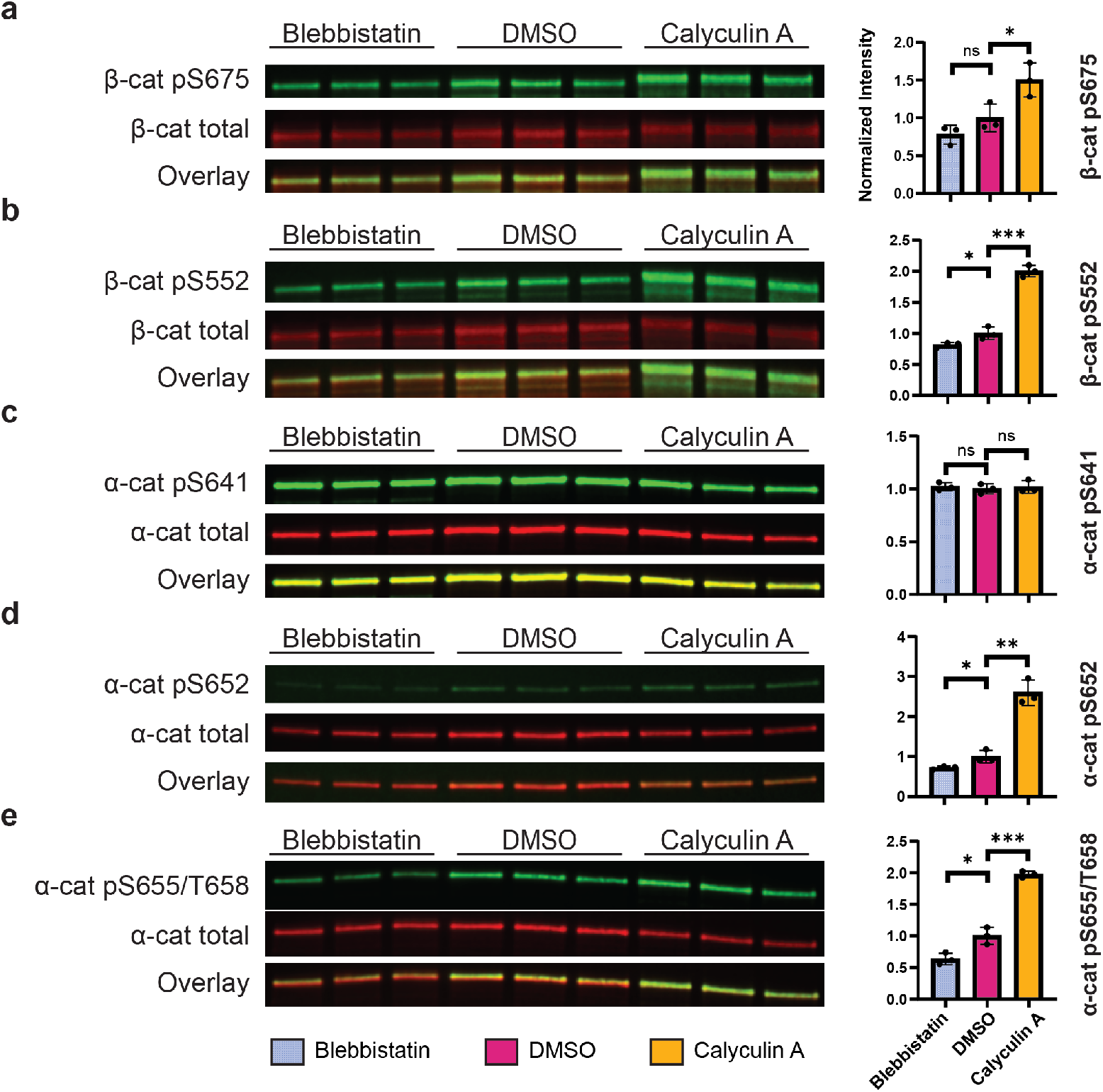
Western blots of cadherin-associated β- and α-catenin purified from HEK cells treated with blebbistatin, DMSO (control) and calyculin A. (a-b) β-cat S675 and S552 are mechano-sensitive. (c) α-cat S641 phosphorylation is sustained and independent of actomyosin perturbation. (d-e) α-cat S652, S655 and T658 are actomyosin-sensitive. All experiments were performed in triplicates. Intensities of catenin phospho-antibodies were divided by corresponding total catenin intensities and normalized to the ratio in DMSO controls. Error bars: one standard deviation. ns: *P>*0.05; *: *P*<0.05; **: *P*<0.01; ***: *P*<0.001 (Student’s *t*-test).

The proteoform landscapes of β-cat (**Figure 3b**) suggested the presence of up to 10 phosphosites, whereas only 9 were identified in our BUP analysis. The DMSO condition revealed evidence of a third phosphorylation relevant to adhesion that may have been missed. Indeed, less-abundant sites such as S102 and Y654, reported in the literature to be associated with adherens junctions, may contribute. Notably, Y654 has been shown to be mechano-sensitive, where its phosphorylation reduces β-cat binding affinity to E-cad and disrupts the complexes.^[46, 57]^ This justifies the absence of pY654 in the Ecad-cyto complexes. On the other hand, S102, together with S29 (identified in this study) and T112, was previously identified as a substrate of casein kinase II (CKII) that promotes binding of β-cat with α-cat.^[58]^ While we detected the unphosphorylated peptide corresponding to S102 (**Figure S6a**), we cannot rule out the presence of pS102 at low stoichiometry (**Table S1b**). A S102 site-specific phospho-antibody is currently not available. This observation further underscores the importance of profiling full proteoform landscapes for accurate identification of modification sites involved in cell-cell adhesion.

The phospho-domain of α-cat (S641–D660), also known as the P-linker, has been associated with adherens junction strength, as mutants that block phosphorylation interfere with dynamic epithelial movements.^[33, 59-60]^ This region contains five well-established phosphorylation sites—S641, S652, T654, S655, and T658—all of which were identified in Ecad-cyto complexes purified from calyculin A-treated cells (**Figure S6b, Table S1c**). Surprisingly, the intensity of pS641 remained consistent across the three treatment conditions (**Figure 4c**). Together with the α-cat proteoform landscapes, which showed no unphosphorylated α-cat (**Figure 3b**), these data suggest that S641 is constitutively phosphorylated within cadherin-catenin complexes. This finding is significant because it challenges the conventional approach of studying the functional consequences of P-linker phosphorylation using all-on/off phospho-mutants—where all phosphosites are mutated to A (phospho-null) or D/E (phospho-mimic)—to assess their effects on cadherin-catenin complex formation or apical junction localization.^[33, 52]^ Our results suggest that an all-off phospho-null α-cat mutant may not reflect the physiological state at cell-cell junctions and could lead to poor correlation with cellular functions or phenotypes. Interestingly, prior studies have demonstrated that phospho-null α-cat mutants are still capable of forming homodimers.^[33]^ This raises the possibility that phosphorylation at S641 favors α-cat participation in junction organization rather than cytosolic dimeric complexes. These observations underscore that proteoforms represent precise molecular entities with distinct primary structures underlying specific cellular functions. Therefore, studying proteins at the proteoform level is essential, rather than focusing solely on individual or aggregated PTM sites.

We further confirmed that the remaining P-linker phosphosites—S652, T654, S655, and T658—are mechano-sensitive, with their phosphorylation levels (as detected by phospho-antibodies) significantly increasing in response to elevated actomyosin contractility (**Figure 4d–e**). It is important to note that pT654, pS655, and pT658 are indistinguishable by antibodies due to their close proximity. Two additional phosphosites in the actin-binding domain of α-cat—S851 and S886—were previously identified only in large-scale phospho-proteomics datasets,^[53]^ and phospho-specific antibodies for these sites are currently unavailable. Their individual functions, such as whether they facilitate α-cat–F-actin binding, and their proteoform-level regulation mechanisms, including potential dependence on P-linker phosphorylation, remain to be investigated in context of recent cryoEM data of the α-cat/F-actin interface.^[61]^ Notably, the total number of phosphosites observed in the α-cat proteoform landscapes (**Figure 3b**) corresponds with the seven phosphosites identified in the BUP experiment.

### New Method Significantly Increased Catenin Sequence Coverage in Top-Down Mass Spectrometry

We recently demonstrated that individual ion mass spectrometry not only facilitates MS^1^-level (intact protein) measurements, but also enables MS^2^-level acquisition of protein fragment ions created by various fragmentation techniques available on Orbitrap instruments. This approach—termed I^2^MS^2^—substantially enhances sequence coverage of large proteoforms (>30 kDa). For example, in the case of MEK1, sequence coverage of the 43 kDa proteoform increased from 3.5% using traditional ensemble measurements to over 50% with I^2^MS^2^.^[23]^ The potential of this method for proteoforms exceeding 50 kDa remains largely unexplored.

To fragment cadherin-associated catenins, Ecad-cyto complexes were ionized under denaturing conditions and introduced into an Orbitrap Eclipse. Precursor ions were isolated using a quadrupole, subjected to either higher-energy collisional dissociation (HCD) or electron transfer dissociation (ETD), and subsequently detected in the Orbitrap at low ion densities. This configuration allowed for the observation of individual ion events and accumulation of signal for charge assignment using slope-based analysis (**Figure 1**). Using HCD in conjunction with I^2^MS^2^, we achieved approximately 25% sequence coverage of β-cat (**Figure S8a**), a substantial improvement over the <1% typically obtained using LC-MS/MS with conventional ensemble measurements (**Figure S8b**). In these traditional methods, limitations such as *m/z* space crowding, ion decay, and low fragmentation efficiency restrict the detection of informative, deconvolutable fragments. Likewise, α-cat sequence coverage was increased to ∼30% using ETD and I^2^MS^2^ (**Figure S9a**), compared to <1% coverage with LC-MS/MS (**Figure S9b**).

While I^2^MS^2^ significantly improves fragment ion detection and sequence coverage of large proteoforms, limitations persist. Specifically, the quadrupole isolates ions based on *m/z* within a narrow range (∼1000 Th for most denatured proteins), which becomes congested in the absence of LC separation. This range may include multiple proteoforms, a distribution of charge states, and overlapping isotopic peaks spaced by <1 Th—differences below the resolution threshold of quadrupole isolation. Consequently, isolating a single catenin proteoform for unambiguous structural characterization remains challenging. Emerging strategies, such as proton transfer charge reduction (PTCR)-assisted I^2^MS^2^,^[24]^ may alleviate these precursor selection issues. Nevertheless, our current I^2^MS^2^ data confirm that α-cat S641 is constitutively phosphorylated (**Figure S9a**), representing a significant advancement in our understanding of α-cat phospho-regulation at the proteoform level—enabled by the unique capabilities of the I^2^MS platform.

### A Phospho-Code for Cadherin-Catenin Complex Regulation

The rapid regulation of cell–cell cohesion in response to mechanical force is critical for maintaining epithelial barrier integrity under dynamic tissue conditions. This has led to considerable interest in the mechano-sensing properties of adhesive proteins.^[62]^ Our data suggest that catenins—key components of adherens junctions—are mechano-sensitive, and their phosphorylation patterns, or phospho-proteoforms, are associated with the levels of actomyosin tension they experience. Drawing a parallel to the “histone code,”^[63-64]^ wherein combinations of complex methylation, acetylation, phosphorylation and other PTMs on histone proteoforms regulate gene expression, we propose a “catenin phospho-code” hypothesis for adherens junction organization (**Figure 5**). In this model, cadherin–catenin function is directed by distinct catenin proteoforms that are chemically modified in response to actomyosin tension, thereby determining junctional architecture.

**Figure 5.**
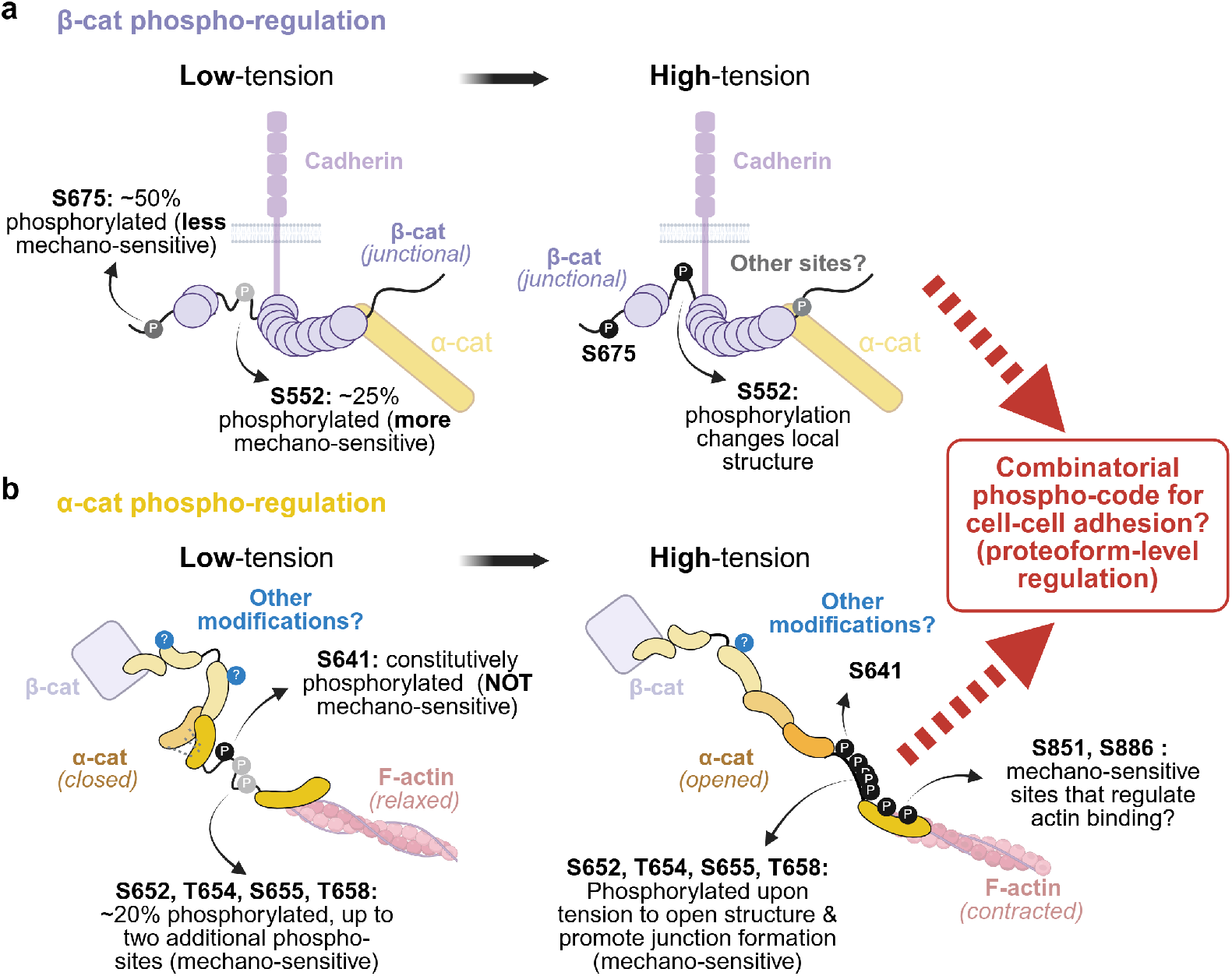
Proposed β- and α-catenin actomyosin tension/phospho-regulation models. (a) β-cat S675 and S552 are both regulated by actomyosin activity, with S552 being more sensitive. It is possible that there are additional sites involved such as S102, observed by others but not captured by this study. (b) α-cat S641 is constitutively modified, whereas S652, T654, S655, T658 in the P-linker region are actomyosin sensitive. We also identify two new actomyosin-sensitive sites (S851 and S886) in the actin-binding domain. This work implies that these sites may work in concert for proteoform-level regulation of adherens junction organization. Although phosphorylation is the major modification of α-cat, there may also be other unknown modifications involved.

In this study, we demonstrate that both β- and α-catenins are mechano-sensitive through their phospho-proteoforms. For β-cat, the primary mechano-sensitive sites are S675 and S552 (**Figure 5a**). Under low tension, approximately 50% of S675 and 25% of S552 are phosphorylated, suggesting that S552 is more responsive to increased actomyosin force. We also discuss potential additional phosphosites within β-cat’s α-cat-binding domain, including S102 and T112. In α-cat, we find that S641 is constitutively phosphorylated, whereas other residues within the P-linker domain are mechano-sensitive and exhibit hyperphosphorylation under high-tension conditions, like mitosis (**Figure 5b**).^[29]^ Furthermore, our data suggest that two newly identified phosphosites within the actin-binding domain (S851 and S886) may also be mechano-sensitive and could influence α-cat’s interaction with actin filaments, although their interplay with P-linker modifications remains uncharacterized. Our top-down proteomics approach provides new insights into how multiple phosphorylation sites on catenins may act cooperatively to regulate adherens junction organization.

## Conclusion

We have extended the capability of proteoform characterization beyond 100 kDa using individual ion mass spectrometry, demonstrating that cadherin-associated catenins are regulated at proteoform-level in response to actomyosin force disruptors. Our analysis revealed phosphorylation as the major modification on catenins and identified both mechano-sensitive phosphosites and the constitutively modified residue pS641 on α-catenin within cadherin-catenin complexes. Based on these findings, we propose a catenin phospho-code hypothesis for adherens junction organization. This work establishes a foundation for proteoform-level insights into the regulation of cell–cell adhesion and has improved top-down proteomics.

## Supporting Information

The authors have cited additional references within the Supporting Information.^[65-66]^

## Acknowledgements

This work was supported in part by the National Institute of General Medical Sciences of the National Institutes of Health under Award Number RM1GM156535 to N.L.K. and National Heart, Lung, and Blood Institute under Award Number R01 GM129312, HL163611 to C.J.G. Bottom-up proteomics services were performed by the Northwestern Proteomics Core Facility, generously supported by NCI CCSG P30 CA060553 awarded to the Robert H Lurie Comprehensive Cancer Center and instrumentation award (S10OD025194) from NIH Office of Director. Figures 1, 2, 5 and TOC were created in https://BioRender.com with publication license.

## Note

N.L.K. is involved in entrepreneurial activities in top-down proteomics and consults for Thermo Fisher Scientific. The other authors have declared that no conflict of interest exists.

## Entry for the Table of Contents

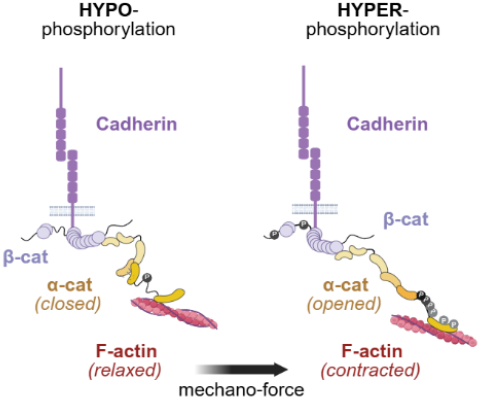

This work profiles cadherin-associated catenin phospho-proteoforms under different actomyosin tensions using top-down mass spectrometry.

