## Supplementary material for "Top-Down Individual Ion Mass Spectrometry Reveals 85-110 kDa Catenin Phospho-Proteoforms Regulated by Actomyosin Contractility": SI

[a] C.-F. Huang, T. Su, N. L. Kelleher*

Northwestern University

[b] A. S. Flozak, C. J. Gottardi*

#### Table of Contents

**Figure S1** $I^2MS$ spectrum of GST-β-catenin ....................S-3
**Figure S2** The proteoform landscape of β-catenin ....................S-4
**Figure S3** The proteoform landscape of α-catenin ....................S-5
**Figure S4** Junctional enrichment of catenins is actomyosin-dependent ....................S-6
**Figure S5** LC-MS analysis of cadherin-catenin complexes ....................S-7
**Figure S6** Bottom-up analysis of calyculin A-treated Ecad-cyto complexes ....................S-8
**Figure S7** β-cat N-terminal sites ....................S-9
**Figure S8** Top-down fragmentation of β-catenin ....................S-10
**Figure S9** Top-down fragmentation of α-catenin ....................S-11
**Experimental Section**....................S-12
**Full-Length Western Blots and Gels**....................S-16
**Additional References** ....................S-18
**Table S1** FragPipe Search Results…………………..xlsx

### Figure S1

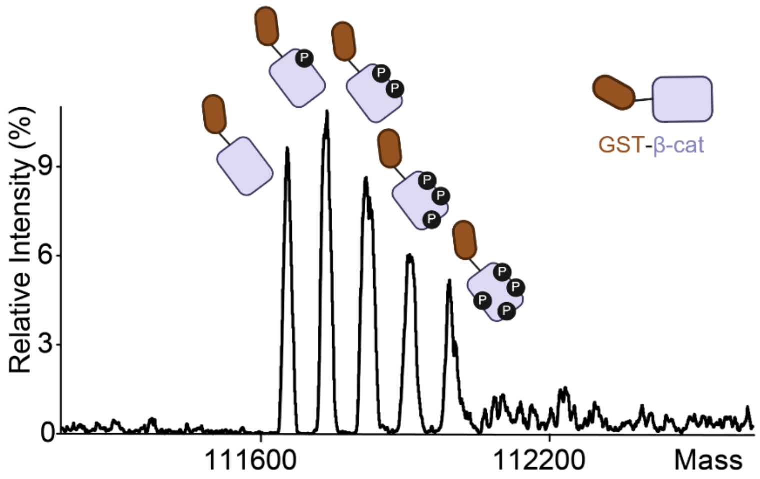


| Proteoform | Observed Mass (Da) | Δm to the previous peak (Da) |
|---|---|---|
| GST-β-cat (M) | 111657.6 | - |
| M+1Phos | 111736.9 | 79.3 |
| M+2Phos | 111816.4 | 79.6 |
| M+3Phos | 111897.3 | 80.9 |
| M+4Phos | 111977.5 | 80.2 |

**Figure S1.** $I^2$MS spectrum of GST-β-catenin. Recombinant GST-fused β-catenin (Abcam ab63175, ~112 kDa) was profiled with optimized $I^2$MS parameters in denatured mode, revealing proteoforms separated by 80 ± 1 Da corresponding to phosphorylation. Up to four phosphorylations were observed.

### Figure S2

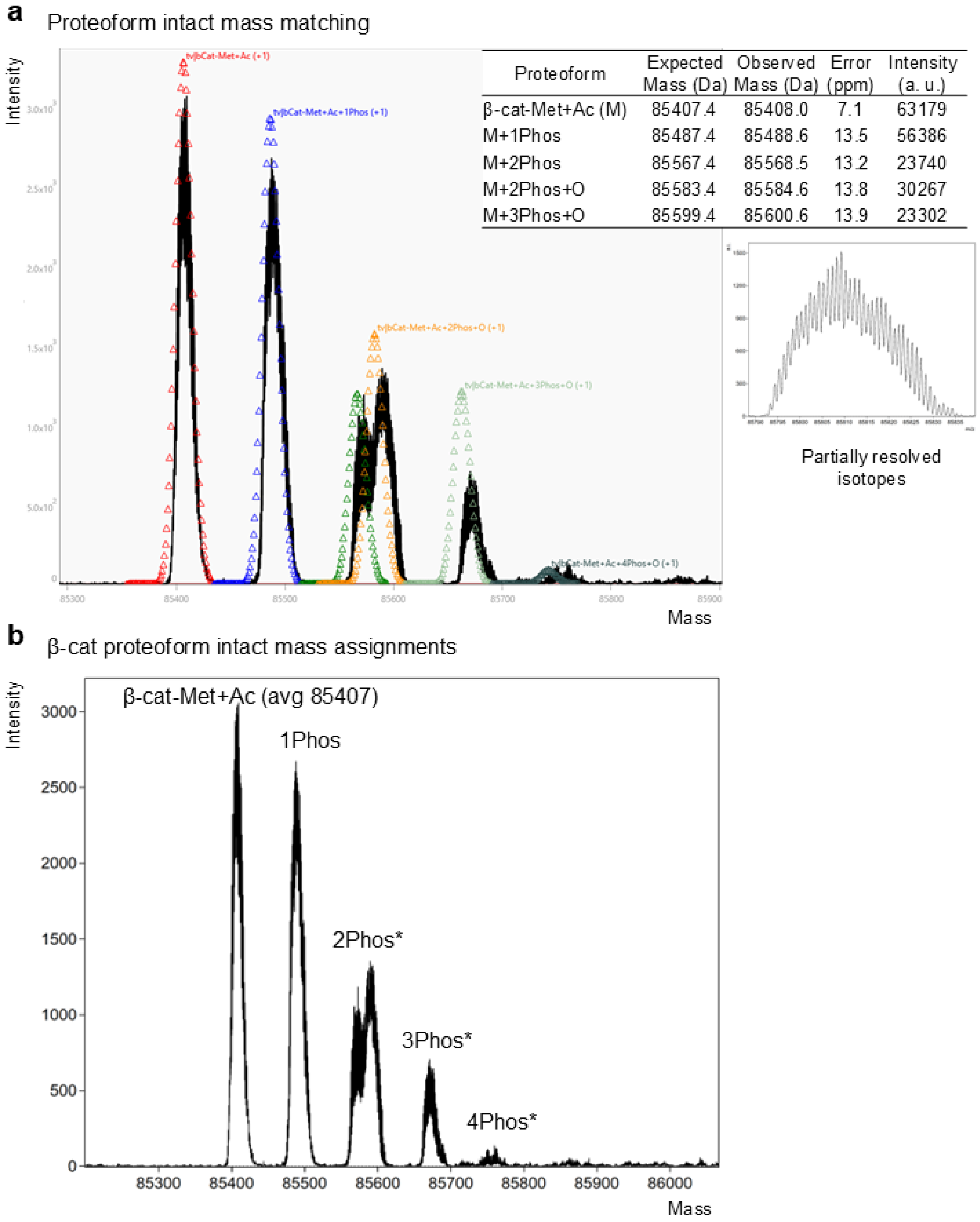


| Proteoform | Expected Mass (Da) | Observed Mass (Da) | Error (ppm) | Intensity (a. u.) |
|---|---|---|---|---|
| β-cat-Met+Ac (M) | 85407.4 | 85408.0 | 7.1 | 63179 |
| M+1Phos | 85487.4 | 85488.6 | 13.5 | 56386 |
| M+2Phos | 85567.4 | 85568.5 | 13.2 | 23740 |
| M+2Phos+O | 85583.4 | 85584.6 | 13.8 | 30267 |
| M+3Phos+O | 85599.4 | 85600.6 | 13.9 | 23302 |

**Figure S2.** The proteoform landscape of β-catenin. (a) Intact mass matching considering phosphorylation and oxidation. Isotopic patterns are partially resolved. (b) β-cat proteoform intact mass assignments. Up to three phosphorylations are observed. *: including oxidized proteoforms or phosphate adducts.

#### Figure S3

**a** Proteoform intact mass matching

| Proteoform | Expected Mass (Da) | Observed Mass (Da) | Error (ppm) | Intensity (a. u.) |
|---|---|---|---|---|
| α-cat-Met+Ac (M) | 99982.1 | Not Observed | | |
| M+1Phos | 100062.1 | 100060.8 | -13.3 | 23653 |
| M+2Phos | 100142.1 | 100142.7 | 6.2 | 6224 |
| M+2Phos+O | 100158.1 | 100157.8 | -2.3 | 9040 |
| M+3Phos | 100222.1 | 100221.8 | -2.8 | 3857 |
| M+3Phos+O | 100238.1 | 100236.7 | -13.7 | 6052 |
| M+3Phos+2O | 100254.1 | 100252.3 | -17.9 | 3546 |

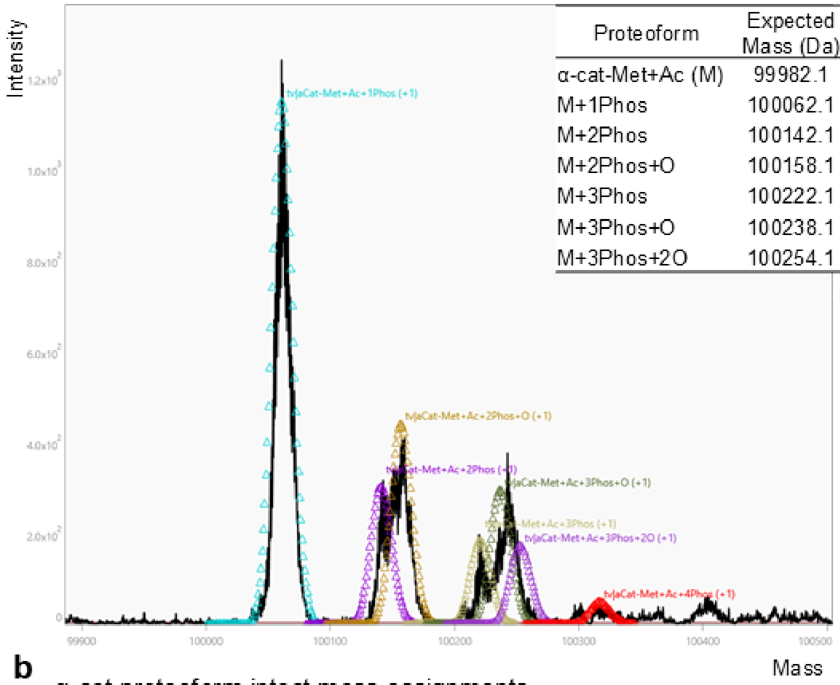


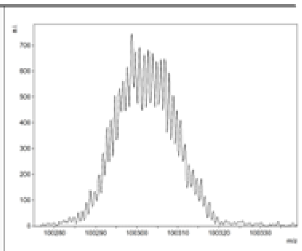


**b** α-cat proteoform intact mass assignments

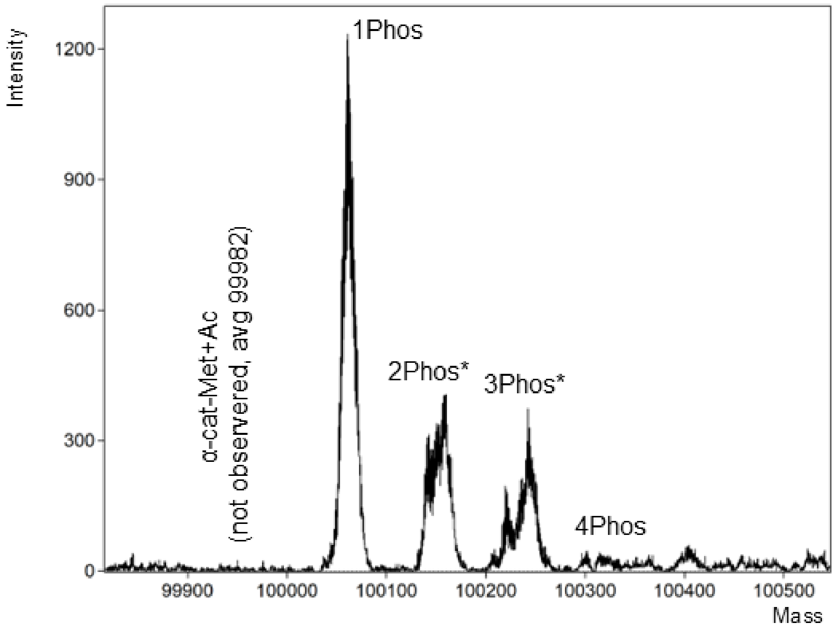


**Figure S3.** The proteoform landscape of α-catenin. (a) Intact mass matching considering phosphorylation and oxidation. Isotopic patterns are partially resolved. (b) α-cat proteoform intact mass assignments. There is no unphosphorylated α-cat. There is no unphosphorylated α-cat detected in the cadherin complex. One to three phosphorylations are observed. *: including oxidized proteoforms or phosphate adducts.

#### Figure S4

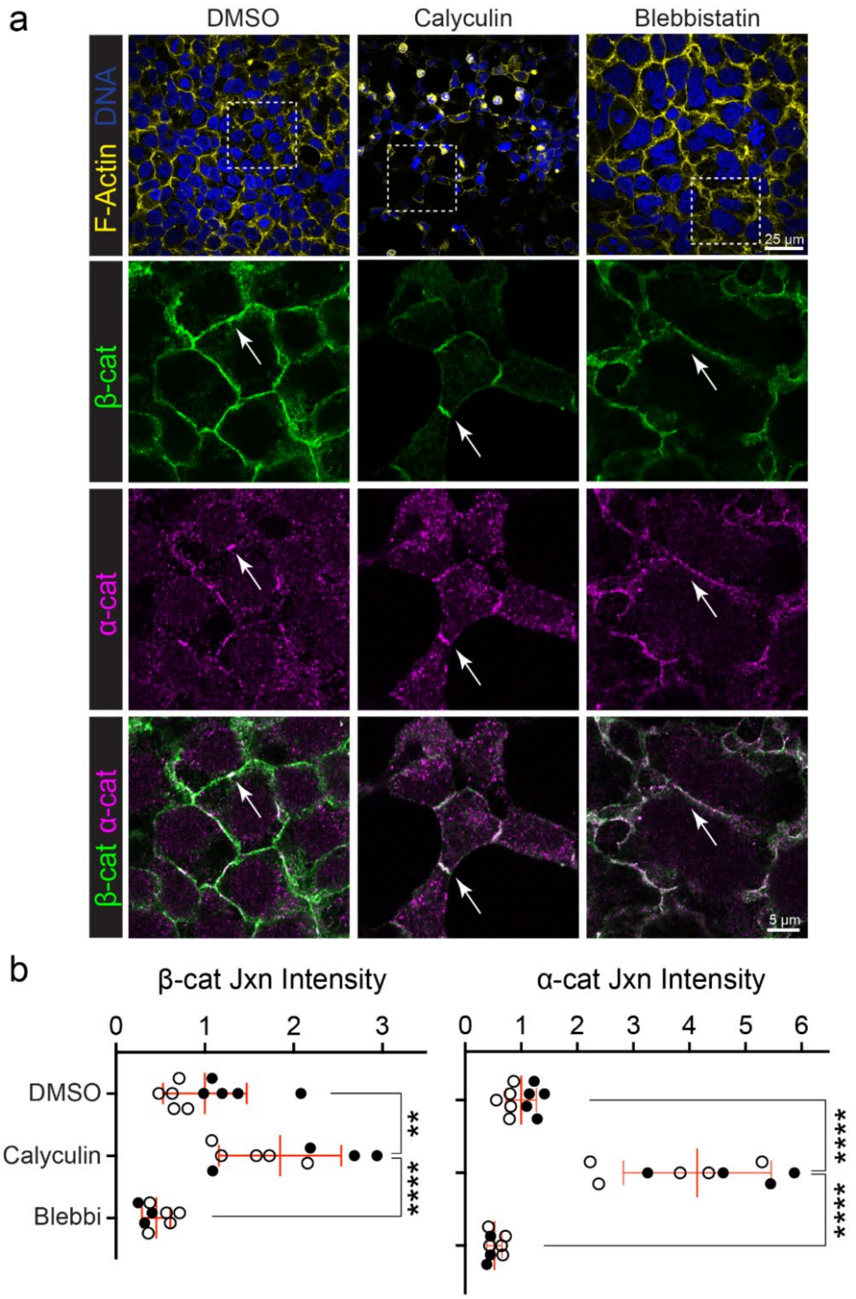


**Figure S4.** Junctional enrichment of catenins is actomyosin-dependent. (a) HEK cells treated with DMSO (control), calyculin (actomyosin activator), and blebbistatin (actomyosin inhibitor) were stained for DNA (blue), F-actin (yellow), β-cat (green) and α-cat (magenta) under a fluorescent microscope. Arrow: adherens junction. (b) Graphs show β-cat and α-cat fluorescence intensities at the junction (Jxn minus extra-junctional cytoplasmic intensity) under different actomyosin-perturbing conditions. Calyculin increased catenin junction enrichment whereas blebbistatin decreased enrichment. Error bars: one standard deviation. **: $P<0.01$; ****: $P<0.0001$ (Student's *t*-test).

#### Figure S5

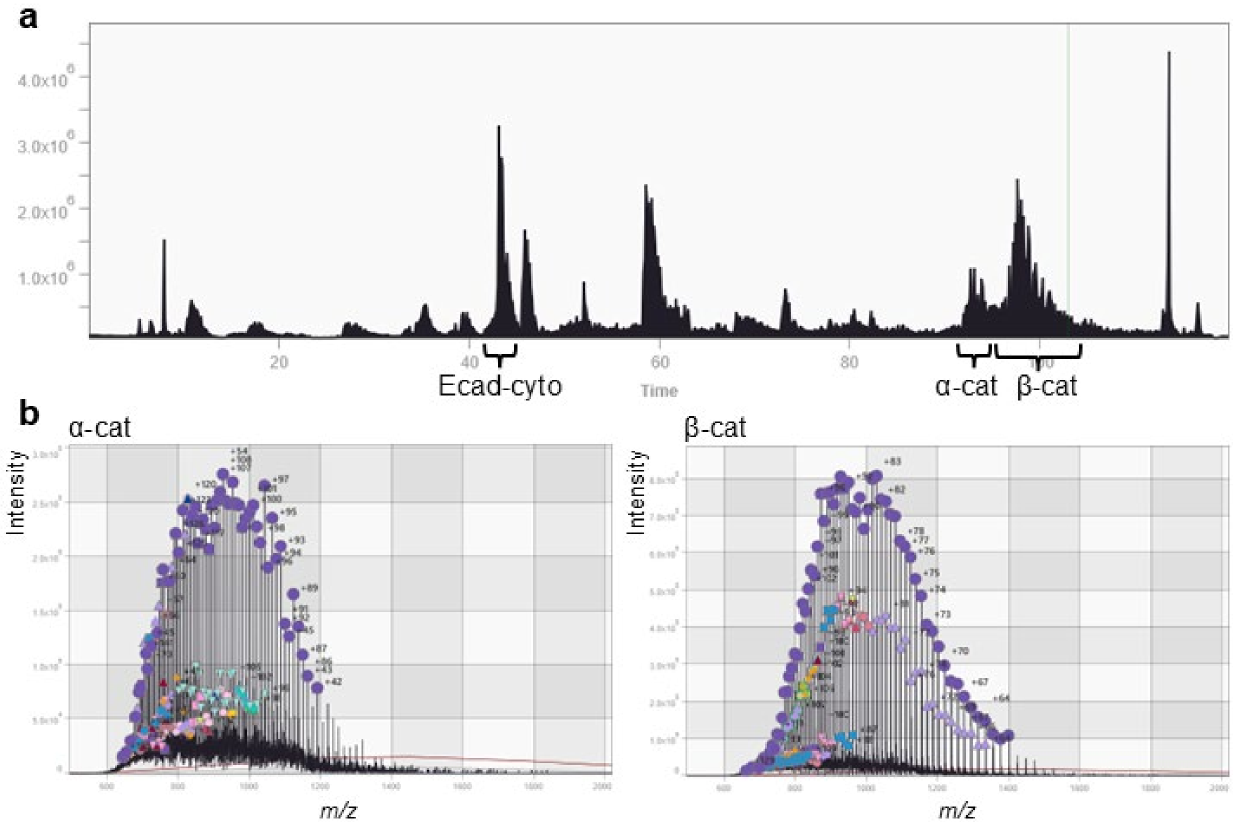


**Figure S5.** LC-MS analysis of cadherin-catenin complexes. (a) Chromatogram of Ecad-cyto eluate analyzed over a 2 h gradient. (b) Ensemble MS is low resolution for highly charged catenins, resulting in extensive spectral congestion of proteoforms.

#### Figure S6

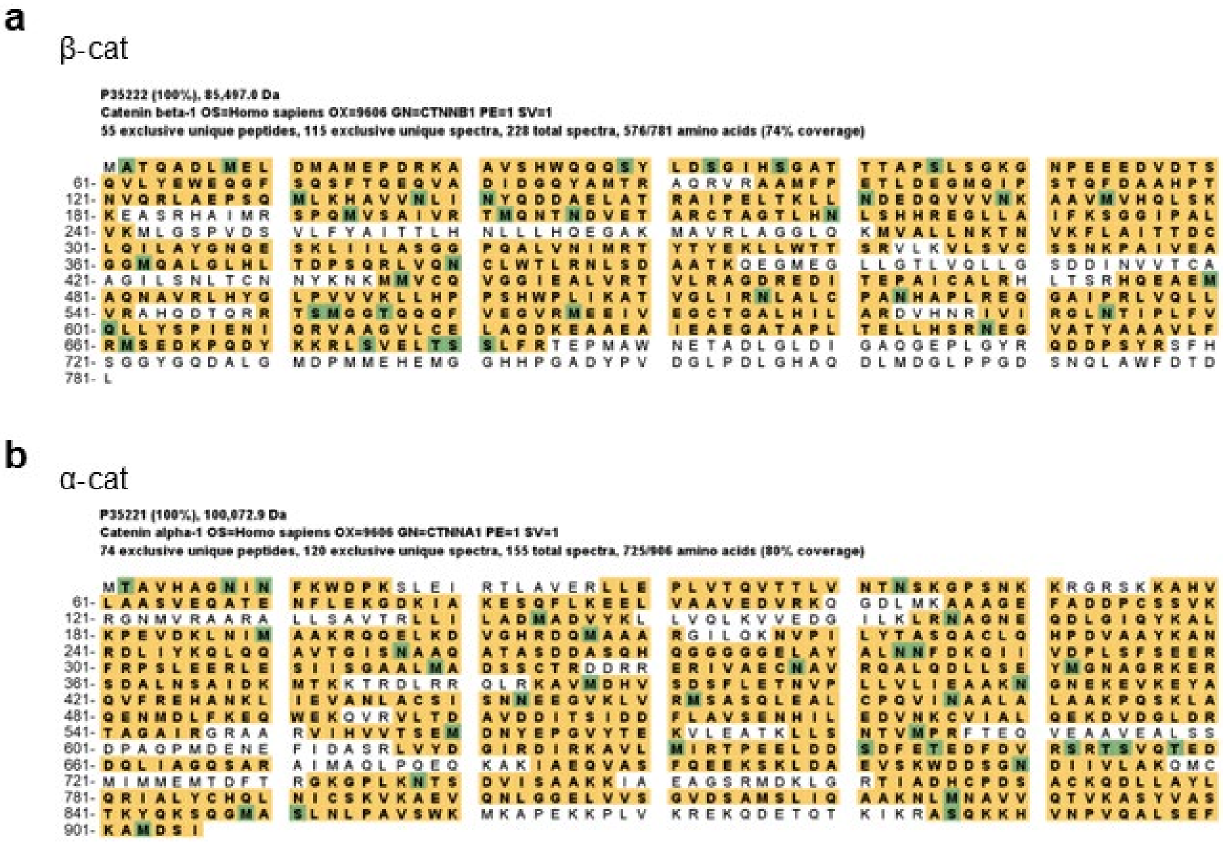


**Figure S6.** Bottom-up analysis of calyculin A-treated Ecad-cyto complexes. (a) Sequence coverage map of β-cat. (b) Sequence coverage map of α-cat. Peptide search was performed on MASCOT including variable phosphorylation at S, T, Y residues and the results were visualized with Scaffold software. Yellow: identified peptides; green: modified residues (including ambiguous modifications).

#### Figure S7

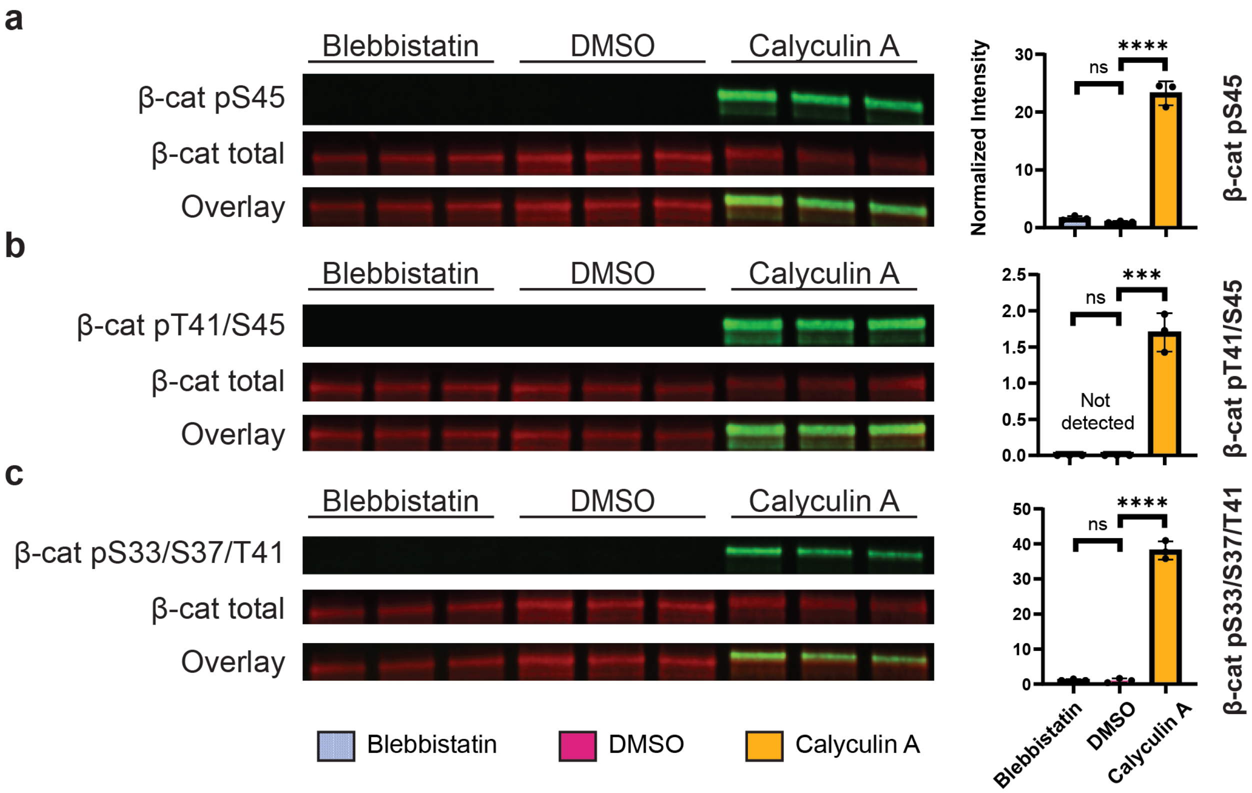


**Figure S7.** β-cat N-terminal sites. (a-c) pS45, pT41, pS37 and pS33 are only detected in HEK cells treated with calyculin A. These sites are traditionally associated with GSK3β-mediated degradation pathways. All experiments were performed in triplicates. Intensities of catenin phospho-antibodies were divided by corresponding total catenin intensities and normalized to the ratio in DMSO controls. Error bars: one standard deviation. ns: $P$>0.05; *: $P$<0.05; **: $P$<0.01; ***: $P$<0.001; ****: $P$<0.0001 (Student's $t$-test).

#### Figure S8

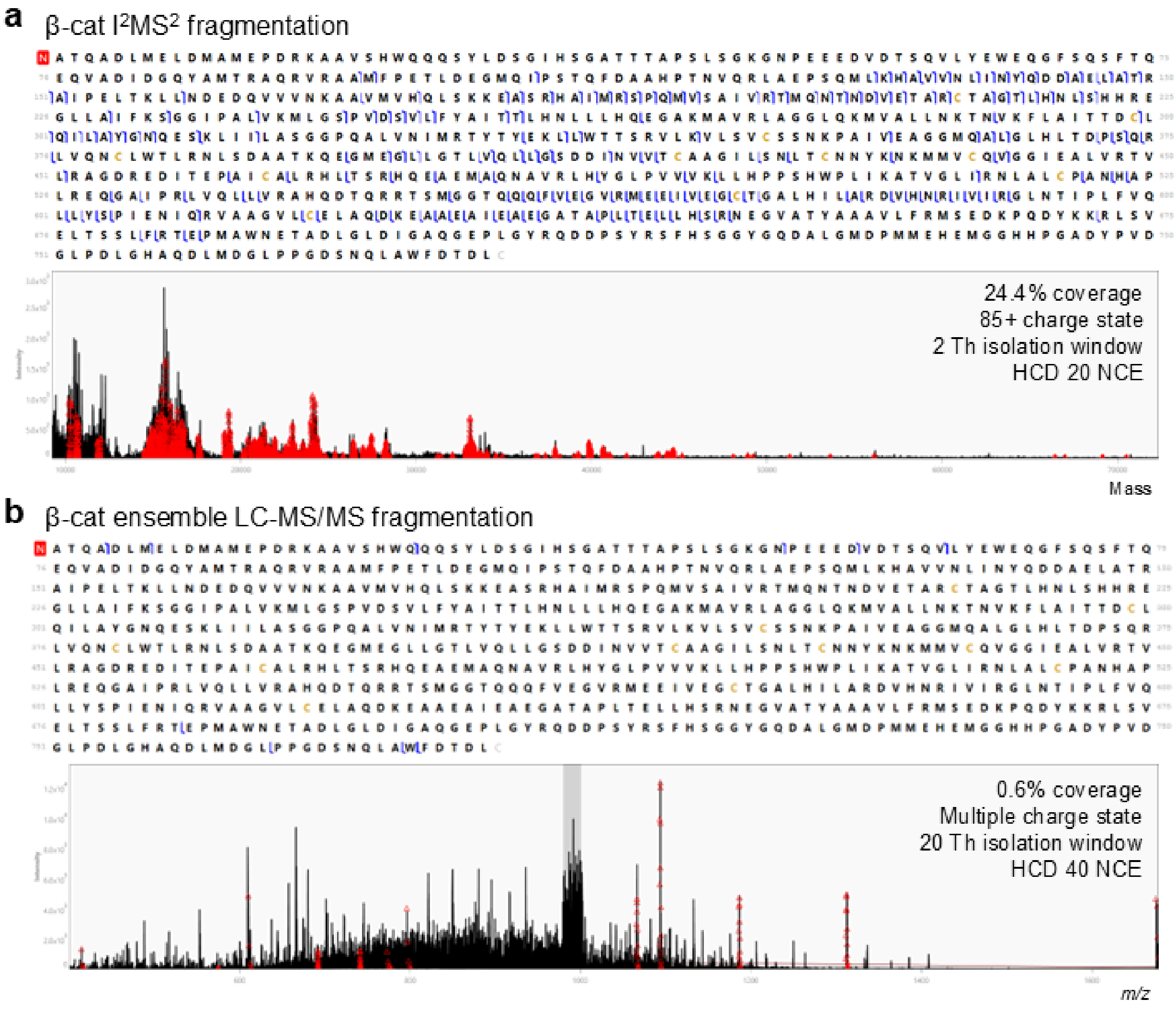


**Figure S8.** Top-down fragmentation of β-catenin. (a) β-cat fragment ion detection was significantly improved by individual ion mass spectrometry for $MS^2$-level experiments. (b) Representative β-cat LC-MS/MS spectrum and sequence coverage map acquired using traditional ensemble detection method.

### Figure S9

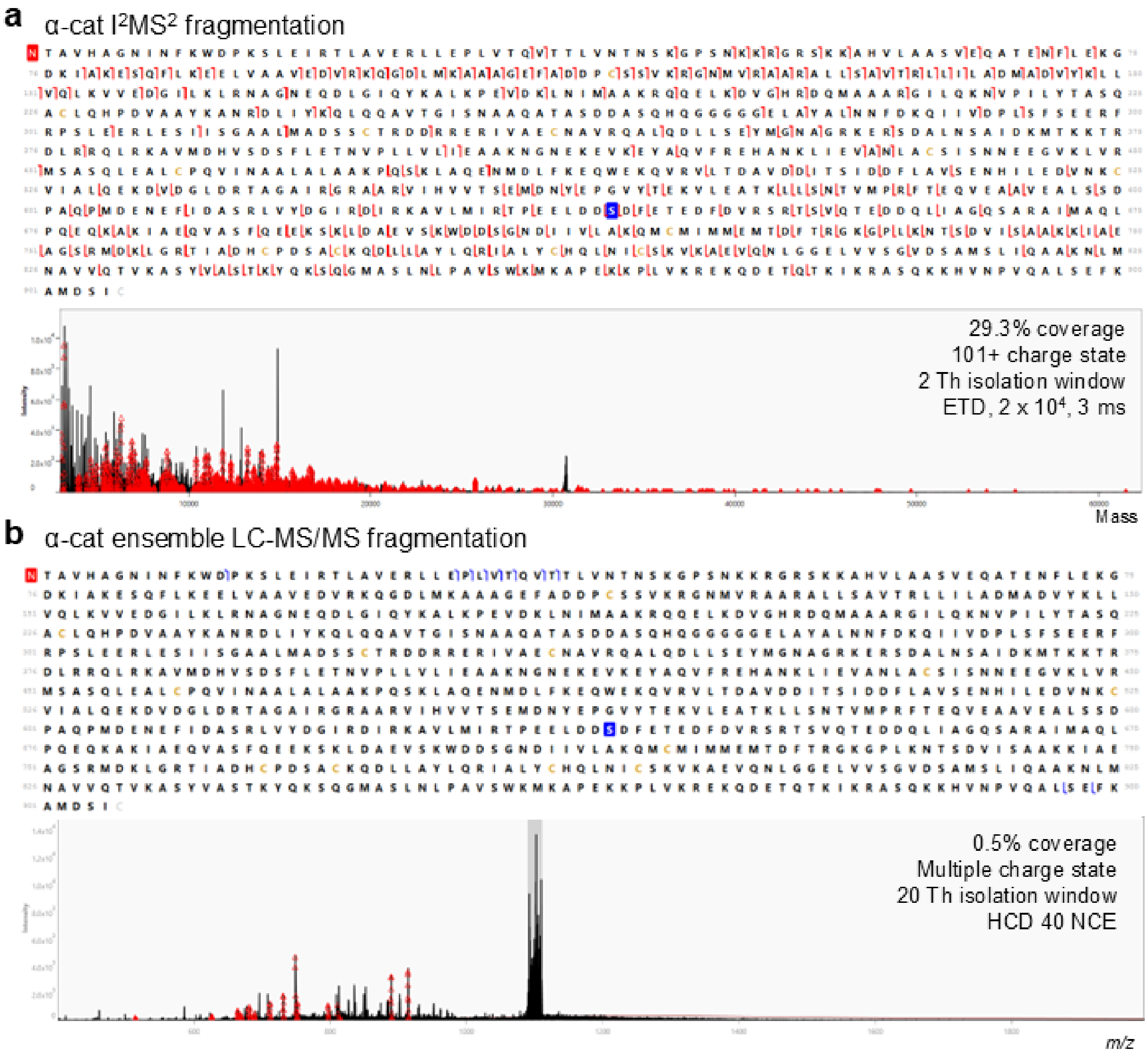


**Figure S9.** Top-down fragmentation of α-catenin. (a) α-cat fragment ion detection was significantly improved by individual ion mass spectrometry for $MS^2$-level experiments. (b) Representative α-cat LC-MS/MS spectrum and sequence coverage map acquired using traditional ensemble detection method.

#### Experimental Section

**Antibodies.** The following primary antibodies were used in this study for western blots: mouse anti-β-catenin (610153, BD Bioscience), mouse anti-α-catenin (ALX-804-101, Enzo), rabbit anti-α-catenin (3236, Cell Signaling), mouse anti-c-myc (M4439, Millipore Sigma), rabbit anti-β-catenin pS33/S37/T41 (9561, Cell Signaling), rabbit anti-β-catenin pT41/S45 (9565, Cell Signaling), rabbit anti-β-catenin pS45 (9564, Cell Signaling), rabbit anti-β-catenin pS552 (9566, Cell Signaling), rabbit anti-β-catenin pS657 (9567, Cell Signaling), rabbit anti-α-catenin pS641 (11330, Signalway), rabbit anti-α-catenin pS652 (13061, Cell Signaling), and rabbit anti-α-catenin pS655/T658 (13231, Cell Signaling). The following primary antibodies were used in this study for immunofluorescence and microscopy: mouse anti-β-catenin (610153, BD Bioscience), rabbit anti-α-catenin (3236, Cell Signaling), Alexa Fluor 568 phalloidin (A12380, Invitrogen), Alexa Fluor 647 phalloidin (A22287, Invitrogen). The secondary antibodies were: IRDye 680 RD donkey anti-mouse (926-68072, LI-COR, western blot), IRDye 800 CW donkey anti-rabbit (926-32213, LI-COR, western blot), Alexa Fluor 488 goat anti-mouse (A-11001, Invitrogen, microscopy), and Alexa Fluor 568 goat anti-rabbit (A-11001, Invitrogen, microscopy).

**Cell culture.** HEK 293T cells were obtained from American Type Culture Collection (ATCC). The Ecad-cyto HEK 293T cell line was previously created and validated.[1] The plasmid sequence was recently checked with Sanger sequencing for authentication. Cells were maintained in Dulbecco's Modified Eagle's Medium (DMEM, Corning), containing 10% fetal bovine serum (FBS, Atlanta Biologicals), 100 U/ml penicillin and 100 mg/mL streptomycin (Corning) at 37°C and 5% $CO_2$. Ecad-cyto HEK cells were supplemented with 0.25 mg/mL Hygromycin B (Gibco). For actomyosin treatments, blebbistatin (10 mM, Millipore Sigma) and calyculin A (10 μM, Millipore Sigma) stock solutions were prepared in sterile DMSO. Cells were treated with blebbistatin (20 μM, 24 h), DMSO (control, 0.2%, 24 h) or calyculin A (20 nM, 2 h) in fresh media.

**Confocal microscopy.** Microscope cover glasses (22 x 22 mm, #1.5, Fisher Scientific) were flamed and coated with 1 mL poly-D-lysine (0.1 mg/mL, Gibco) in a 12 well plate for 1 h at room temperature. HEK cells were seeded at a seeding density of 2.5 x $10^4$ cells and incubated for 4 days and treated with blebbistatin, DMSO or calyculin A in fresh media. Cells were fixed in 4% paraformaldehyde (PFA) in PBS for 15 min, quenched with 100 mM glycine, permeabilized with 0.3% Triton X-100, and blocked with 3% normal goat serum (Millipore Sigma). Primary (1:100) antibodies were incubated at room temperature for 1 h and secondary antibody (1:300) incubations were performed at room temperature for 30 min Phalloidin (1:50) was added to the secondary antibody incubation for F-actin staining. Hoechst (1:10,000, Thermo Fisher) was incubated for 5 min, interspaced by multiple washes in PBS, and followed by mounting coverslips in ProLong Gold fixative (Life Technologies). Fixed cells were imaged on a Nikon A1 (CAM A1RB) microscope with GaAsP detectors and equipped with a 95B prime Photometrics camera using S Fluor 40x Oil DIC H N2 objective or APO 60x I Oil objective. Confocal *z*-stacks were collected at 0.25 μm step size. Captured images were processed in Fiji software. β- and α-catenin intensities were quantified at cell-cell junctions and statistical analyses were performed in GraphPad Prism.

**Cell lysis and immunoprecipitation.** Ecad-cyto HEK cells were grown to confluency in a 150 mm dish (Corning), treated with blebbistatin, DMSO or calyculin A in fresh media, washed with PBS and harvested in cell lysis buffer [50 mM Tris, 150 mM NaCl, 5 mM EDTA, 1% Triton X100, pH 7.5 and 1X Halt protease/phosphatase inhibitor cocktail (Thermo Fisher Scientific)]. Cell lysate was sonicated and centrifuged at 14,000 rpm for 10 min to remove debris. The concentration was determined by Bradford assay (5000006, Bio-Rad). Lysate was incubated overnight with 25 µL myc-trap agarose (ChromoTek) at 4$^{o}$C under agitation. Immunocomplexes were collected by centrifugation and washed once with ice-cold cell lysis buffer, twice with 10X TBS (Fisher Scientific), twice with 1X TBS and once with LC-MS grade water (Fisher Scientific). For TDP experiments, Ecad-cyto complexes were eluted with 100 µL 1% formic acid (LC-MS grade, Fisher Scientific) at 37$^{o}$C for 10 min For BUP and electrophoresis experiments, Ecad-cyto complexes were eluted with 100 µL 2X SDS-PAGE sample buffer (Bio-Rad) containing 5% β-mercaptoethanol (Sigma-Aldrich) at 95$^{o}$C for 10 min.

**Electrophoresis and immunoblotting.** Lysates or IP eluates were resolved by 4-20% Criterion TGX Midi gels (Bio-Rad, 150 V, 90 min) and transferred to nitrocellulose membranes using the Trans-Blot Turbo system (Bio-Rad) following the manufacturer's protocol. Membranes were blocked in Intercept (TBS) blocking buffer (LI-COR) for 1 h at room temperature, incubated overnight with primary antibodies at 4$^{o}$C and then with fluorophore-conjugated secondary antibodies (LI-COR) for 1 h at room temperature. All antibodies were used at the dilutions suggested by the manufacturers. The bands were visualized by LI-COR Odyssey Fc Imaging System. The images were processed with ImageJ and statistical analyses were performed in GraphPad Prism.

**Individual ion mass spectrometry.** IP eluates were buffer exchanged into electrospray-compatible system buffer (69.8% water, 30% acetonitrile, 0.2% formic acid) using the SampleStream platform.[2] Direct injection and I$^2$MS data acquisition was automated using SampleStream. Buffer exchanged sample (70 µL) was picked up by a PAL3 robot, delivered to an Q Exactive HF mass spectrometer (Thermo Fisher Scientific) at a flow rate of 1.2 µL/min, and sprayed through a customed nanospray source. Upon sample injection, a process coined Automated Ion Control (AIC) attenuated ensemble ion signals down to an ion injection time (1–400 ms) corresponding to optimal individual ion collection on a per-sample basis to maximize the number of individual ion signals without producing a large number of multiple ion events at singular *m/z* values.[3] Other instrument parameters included: eFT, off; Orbitrap central electrode voltage, 1 kV; trapping gas pressure, 0.5; spray voltage, 2.6 kV; sheath gas, 0 L/min; in-source CID, 15 eV; source temperature, 320°C; *m/z* acquisition range, 500–3000 *m/z*; resolution 45,000 @ 200 *m/z* (1 s transient time); the data was collected for 1 h. I$^2$MS$^2$ data acquisition was performed using manual direct injection via a syringe pump (1 µL/min) on an Orbitrap Eclipse mass spectrometer (Thermo Fisher Scientific). Precursor ions were selected using a quadrupole with a 2 Th isolation window (centers of *m/z*: β-cat 1011.67, α-cat 991.73). HCD (NCE 20 @ 85+ charge state) was used for β-cat fragmentation and ETD (reagent density 2 x $10^4$, 3 ms reaction time) was used for α-cat. The Orbitrap resolution was 240,000 @ 200 *m/z* (2 s transient time), and the scan range was 400–2,000 *m/z*. The injection time was manually optimized for maximum single ion events (typically 10-50 ms), and the normalized AGC target was 500%. The ion routing multipole (IRM) pressure was set to 0.001 Torr to extend the survival of large fragment ions in $I^2MS^2$ readout. The $I^2MS$ data were acquired for 1 h.

**Data processing.** All of the .storx files collected from $I^2MS$ and $I^2MS^2$ experiments were processed using STORIboard (https://www.proteinaceous.net/storiboard).[4] Charges were assigned with the voting charge assignment algorithm. Based on the ion charge and *m/z*, the accurate mass of each ion was calculated, and a spectrum with all of the charge-assigned ions presenting in the mass domain was generated in a *.mzML* file. Further annotation of $I^2MS$ and $I^2MS^2$ data was performed using TDValidator (Proteinaceous, https://www.proteinaceous.net/tdvalidator).[5] For $I^2MS$, the catenin proteoforms were assigned to spectral peaks according to the matches of intact masses at a maximum tolerance of ± 20 ppm and a mass tolerance of 1 Da. For $I^2MS^2$ data, matches to *b*, *y*, *c* and *z* fragment ions from a specific catenin proteoform were annotated with the following settings: a max tolerance of ± 5 ppm; additional settings in TDValidator were a minimum score of 0.5–0.7 and S/N of 3.

**Top-down LC-MS/MS.** Proteins in the IP eluate were separated using a Vanquish Neo UHPLC chromatographic system (Thermo Fisher). Reversed-phase LC was performed on a MAbPac EASY-Spray column (150 mm length by 150 µm inner diameter, Thermo Fisher Scientific) with an in-house packed PLRP-S trap (25 mm length by 150 µm i.d., Agilent). The total run time was 120 min using a gradient of mobile phase A (99.9% water and 0.1% formic acid) and mobile phase B (19.9% water, 80% acetonitrile, and 0.1% formic acid). The flow rate was set at 1 µL/min, and the gradient used to resolve proteins was 5% B at 0 min, 30% B at 5 min, 55% B at 110 min, 99% B from 111 to 114 min, and 5% B from 115 to 120 min. The column outlet was coupled inline to an EASY-Spray source and an Orbitrap Eclipse or Ascend mass spectrometer (Thermo Fisher Scientific) operating in intact protein mode with 2 mTorr of $N_2$ pressure in the ion routing multiple (IRM). The transfer capillary temperature was set at 320°C, the ion funnel RF was set at 60%, and 15 V of source CID was applied. $MS^1$ spectra were acquired at 7,500 resolving power (at *m*/*z* 200), a normalized AGC target of 1000%, 1200 ms maximum injection time, and 16 µscan. $MS^2$ spectra were acquired in proteoform reaction monitoring (PfRM) mode.[6] Spectra were acquired at 60,000 resolving power (at *m/z* 200), normalized AGC target value of 1200%, 1000 ms of maximum injection time, and 4 µscan/spectrum. Higher-energy collisional dissociation (HCD) was used with the normalized collision energy (NCE) set at 40% to generate $MS^2$ fragmentation spectra. Precursors were quadrupole-isolated by using a 20 Th wide isolation window. The following precursor *m/z* isolation window centers were applied: β-cat 990 *m/z*, α-cat 1100 *m/z*.

**Bottom-up proteomics analysis.** Bottom-up proteomics was performed at Northwestern Proteomics Core Facility. Calyculin A-treated HEK cell Ecad-cyto eluate was digested with trypsin (Promega) following the manufacturer's in-gel digestion protocol. LC-MS/MS analysis was performed on a Vanquish Neo UHPLC (Thermo Fisher Scientific) coupled to an Orbitrap Exploris 240 mass spectrometer (Thermo Fisher Scientific). Analytical separation was conducted using a UHPLC C18 column (Ion Opticks, AUR3-15075C18-CSI). The flow rate was set at 0.2 µL/min Elution of peptides from analytical separation column was performed using a 120 min gradient between buffer A (0.1% formic acid and 99.9% Optima LC/MS grade water) and buffer B (80% acetonitrile, 19.9% Optima LC/MS grade water, and 0.1% formic acid): 0% B at the beginning, 8% B at 1 min, 28% B at 86 min, 50% B at 106 min, 100% B from 107 to 110 min, 0% B from 111 to 120 min. Electrospray ionization using Orbitrap Exploris 240 was performed using a Nanospray Flex Ion Source (Thermo Fisher Scientific) and positive static spray voltage was set at 2400 V. For full $MS^1$, scan range was set to 350-1600 *m/z*. The instrument is set in peptide mode, RF lens: 60%, Orbitrap resolution: 120,000, Normalized AGC Target (%): 300, Maximum injection time (ms): 25, Microscans: 1, and Intensity threshold: 5.0e3. Data-dependent acquisition (DDA) by TopN was performed through fragmentation isolated precursor ions with charges between +2 to +5. The following parameters were used for DDA: Dynamic exclusion mode: exclusion duration (s): 30, mass tolerance: 5 ppm (low) and 5 ppm (high). Data dependent mode: cycle time (s): 2, $ddMS^2$ parameters isolation window (*m/z*): 1.5, Normalized collisional energy: 30%, Orbitrap resolution 15,000, scan range mode: define first mass, first mass (*m/z*): 200, Normalized AGC target (%): 100, Maximum injection time (ms): 50, Microscan: 1. The data was search against the human FASTA database (UniProtProteome: UP000005640) from UniProt using MASCOT[7] and FragPipe with MSFragger for PTM analysis.[8]

**Data availability.** All mass spectrometry *.raw* data, $I^2MS$ STORI files, *.mzML* spectra are available on MassIVE through repository number MSV000098690. There is no password needed to access the files.

#### Full-Length Western Blots and Gels

Fig. 2c

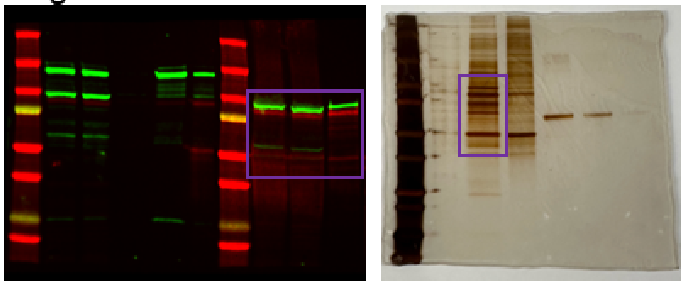

Fig. 4a

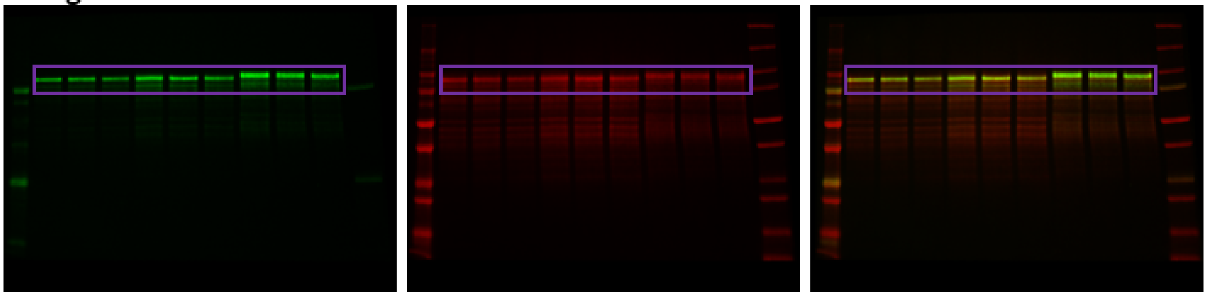

Fig. 4b

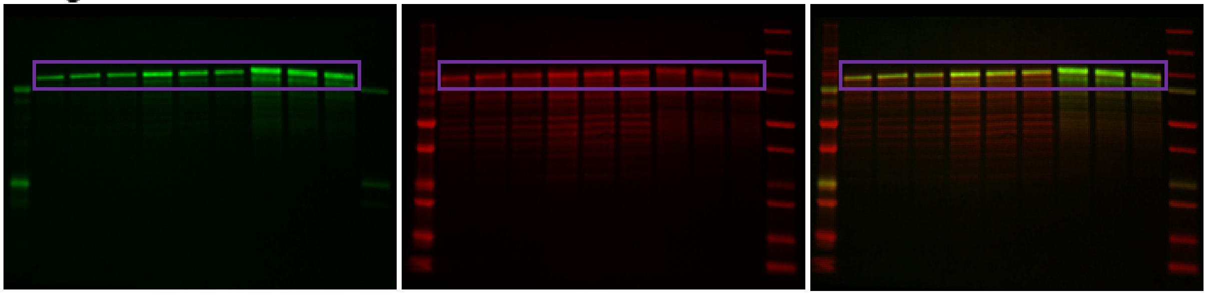

Fig. 4c

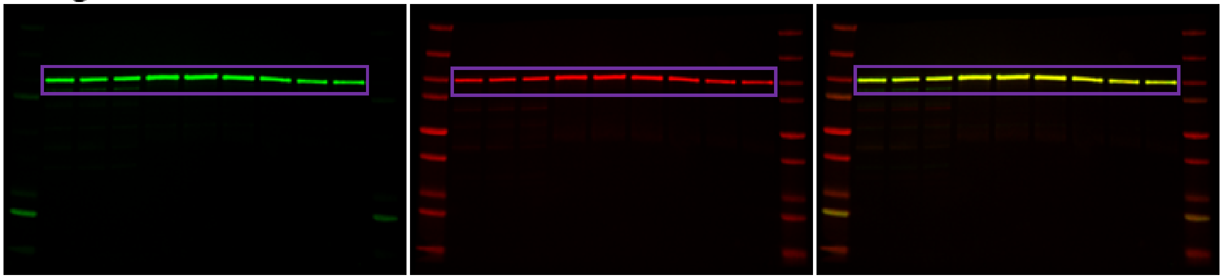

Fig. 4d

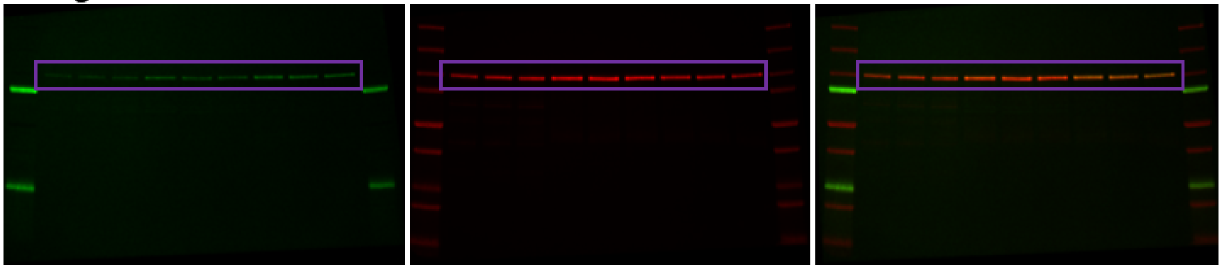

Fig. 4e

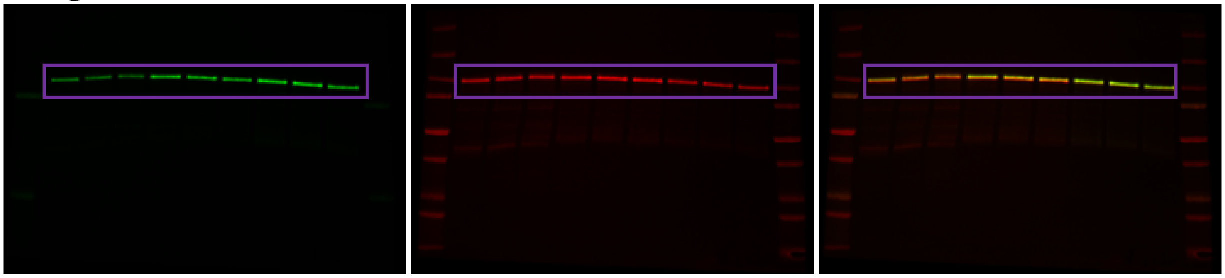

Fig. S7a

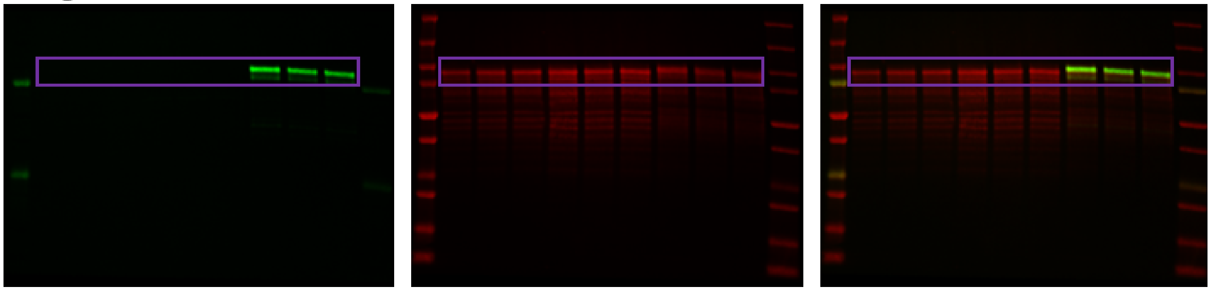

Fig. S7b

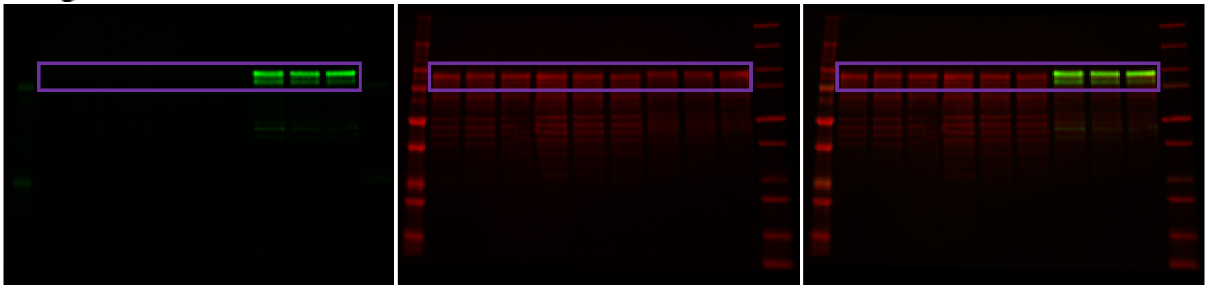

Fig. S7c

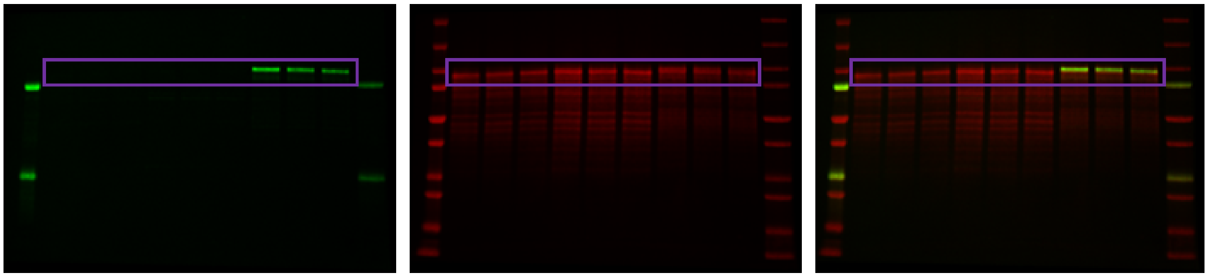
